# Metaproteomics and DNA metabarcoding as tools to assess dietary intake in humans

**DOI:** 10.1101/2024.04.09.588275

**Authors:** Brianna L. Petrone, Alexandria Bartlett, Sharon Jiang, Abigail Korenek, Simina Vintila, Christine Tenekjian, William S. Yancy, Lawrence A. David, Manuel Kleiner

**Affiliations:** Department of Molecular Genetics and Microbiology, Duke University School of Medicine, Durham, NC, United States; Medical Scientist Training Program, Duke University School of Medicine, Durham, NC, United States; Department of Plant and Microbial Biology, North Carolina State University, Raleigh, NC, United States; Duke Lifestyle and Weight Management Center, Durham, NC, United States; Department of Medicine, Duke University School of Medicine, Durham, NC, United States

**Keywords:** gut microbiome, gut microbiota, intestinal microbiome, LC-MS/MS, diet detection, high resolution mass spectrometry, dietary protein sequences, diet database

## Abstract

Objective biomarkers of food intake are a sought-after goal in nutrition research. Most biomarker development to date has focused on metabolites detected in blood, urine, skin or hair, but detection of consumed foods in stool has also been shown to be possible via DNA sequencing. An additional food macromolecule in stool that harbors sequence information is protein. However, the use of protein as an intake biomarker has only been explored to a very limited extent. Here, we evaluate and compare measurement of residual food-derived DNA and protein in stool as potential biomarkers of intake. We performed a pilot study of DNA sequencing-based metabarcoding (FoodSeq) and mass spectrometry-based metaproteomics in five individuals’ stool sampled in short, longitudinal bursts accompanied by detailed diet records (*n*=27 total samples). Dietary data provided by stool DNA, stool protein, and written diet record independently identified a strong within-person dietary signature, identified similar food taxa, and had significantly similar global structure in two of the three pairwise comparisons between measurement techniques (DNA-to-protein and DNA-to-diet record). Metaproteomics identified proteins including myosin, ovalbumin, and beta-lactoglobulin that differentiated food tissue types like beef from dairy and chicken from egg, distinctions that were not possible by DNA alone. Overall, our results lay the groundwork for development of targeted metaproteomic assays for dietary assessment and demonstrate that diverse molecular components of food can be leveraged to study food intake using stool samples.

## Introduction

Biomarkers of food intake are a promising tool for measuring human nutrition, which is the leading preventable risk factor for mortality worldwide.^1^ Food-derived compounds measured in biological specimens can provide objective and accurate data on what is being eaten. Objective measurement strategies can overcome limitations of self-reported dietary data, which are collected by asking individuals to describe what they eat via written record, standardized surveys, or a trained interviewer. While self-reports have provided key insights into the relationship between diet and health, they are at best considered a semi-quantitative strategy with well-characterized sources of error,^2^ including participant memory, cognitive ability, language and cultural suitability of the collection tool, and social desirability bias (*e.g.* a tendency to underreport food intake^3^). Most self-report tools are either administered only once to assess habitual diet for up to the last year (“food frequency questionnaires”), or a handful of times— though there are notable exceptions^4^—to collect repeated data on short-term intake (“24 hour recalls’’). Repeated self-reporting increases the likelihood of reactivity bias, in which individuals alter their diet to make it simpler to report: most existing dietary assessments take between 30 minutes to upwards of an hour to complete, placing both time and cognitive burdens on participants, and depending on the method, research staff. Depending on the specimen type and underlying physiology, biomarkers can facilitate denser sampling strategies to provide high-resolution data on daily variation in diet. Development and validation of dietary biomarkers therefore has the potential to generate more accurate and comprehensive dietary data from large numbers of individuals and to enable retrospective investigations in studies where dietary data were not initially collected.^5^

Prior work on biomarkers of food intake has largely focused on metabolites present in blood, urine, skin, or hair. Stool has been used to a much lesser extent although it is both widely collected in studies and produced directly from consumed foods. Driven by increased research on gut microbiota, large studies now routinely sample stool from hundreds to thousands of individuals.^6–9^ Stool contains molecular information aggregated from host, microbial, dietary, clinical, and environmental factors, and is increasingly leveraged at the population scale with wastewater epidemiology to monitor infectious disease, illicit drug use, or whole-community microbiome composition. After microbial biomass, the largest portion of the dry weight of organic solids in feces is derived from unabsorbed dietary carbohydrate (∼25%), protein (2-25%), and fat (2-15%), with exact proportions varying with the specific foods consumed.^10^ However, much of the biomarker development in stool to date has relied on measuring proxies for residual food material, rather than the food itself. For example, recent efforts have measured fecal metabolite shifts in response to supplementation^11^ or substitution of the source^12^ of dietary protein, or used gut microbiota to predict which of six food items was included in the diet as a controlled intervention.^13^

Direct assessment of food tissue, however, is feasible even after degradation in the gastrointestinal tract. Here, we focus on two macromolecules, DNA and protein, which can provide information on food identity (DNA and protein) and tissue of origin (protein alone). Both can be surveyed with omic-scale tools (“metabarcoding” or “FoodSeq” for DNA, and “metaproteomics” for protein). Dietary DNA has been used to study foods consumed by wild animals, deceased humans (>200 years ago^14^ or recently^15^), and free-living populations^16^. Interestingly, before DNA-based techniques took hold in the field, the earliest animal dietary studies relied on detection of protein from consumed prey tissues. Early work demonstrated that protein epitopes from prey could resist digestion for up to several days.^17^ Metaproteomics, which is the large-scale identification and quantification of proteins from microbiomes using high-resolution mass spectrometry,^18^ has since been applied to query diverse measures in microbial communities^19^ and used to study nutrient flows in biological systems.^20^ Recent work has developed high-throughput methods for stool^21^ that were applied to a human cohort undergoing a dietary intervention.^22^ Plant proteins were observed in the dataset and host and microbial proteomes predicted diet group, but dietary proteins were not systematically identified and analyzed due to lack of a food proteome reference database. This is representative of broader metaproteomics studies of the gut microbiome, which exclude the dietary proteins always contained in the mass spectrometry data by not including them in protein references or removing them as “contaminants.”

To evaluate and compare molecular assessment of dietary proteins and DNA in stool, we applied both metaproteomics and FoodSeq to a pilot set of samples collected in short longitudinal bursts (most in runs of three to five days) from five individuals with detailed accompanying diet records. We began by developing the infrastructure for dietary detection with metaproteomics, which to our knowledge has not yet been applied in humans. To this end, we created the first protein sequence database curated for the identification of dietary peptides and evaluated strategies for analyzing the resulting data. We evaluated broad correspondence between DNA, protein, and diet record measurements, then specifically compared molecular dietary measures to conventional diet records as a reference technique. Finally, we used the high detail of metaproteomics to provide examples of candidate food tissue-specific biomarkers that could be further developed for intake of exact food items.

## Materials and Methods

### Human study design and sample collection

Samples were drawn from two human studies based at Duke University in Durham, NC: a behavioral intervention that returned gut microbiome data to participants (<u>NCT04037306</u>, here “Intervention”) and a protocol for stool collection from healthy donors (Duke Health Institutional Review Board [IRB] Pro00049498, “Habitual Diet”). All participants provided written informed consent and authorized future use of their de-identified stool samples for research. Application of FoodSeq and metaproteomics was a secondary analysis and determined exempt by the Duke Health IRB (Pro00100567).

### Diet detection with metaproteomics

#### Protein extraction and peptide preparation

We homogenized 1.5 g of each sample in 15 mL of SDT-lysis buffer (4% (w/v) SDS, 100 mM Tris-HCl pH 7.6, 0.1 M DTT) with a handheld homogenizer (Omni Micro Homogenizer 115V, Omni International). 1.5 mL of the homogenate was transferred to lysing Matrix E tubes (MP Biomedicals) and bead beaten at 6.45 m/s for 5 cycles of 45s with 1 minute between cycles. Samples were heated to 95°C for 10 minutes, briefly vortexed, and centrifuged at 21,000 x g for 5 minutes to pellet debris. The supernatant was transferred to fresh tubes and centrifuged 3 minutes at 21,000 x g to pellet any remaining debris. We then prepared tryptic digests (13.5 hour digestion) using the filter-aided sample preparation (FASP) protocol^23^ with all centrifugations performed at 14,000 x g. Briefly, we combined 60 uL of the lysate supernatant with 400 uL UA solution (8 M urea in 0.1 M Tris/HCl pH 8.5) in a 10 kDa MWCO 500 µl centrifugal filter unit (VWR International) and centrifuged for 40 minutes. Filters were washed with 200 uL of UA solution and centrifuged for 40 minutes. 100 uL IAA (0.05□M iodoacetamide in UA solution) was added to the filters, incubated 20 minutes, and centrifuged for 20 minutes. Filters were then washed with 100 uL of UA and centrifuged three times, and the buffer changed to 50□mM ammonium bicarbonate by adding 100 µl and centrifuging three times. For digestion, we added 0.95 µg of MS grade trypsin (Thermo Scientific Pierce, Rockford, IL, USA) in 40□µl of 50 mM ammonium bicarbonate to the filters and incubated 13.5 hours in a wet chamber at 37°C. Following digestion, filters were centrifuged for 20 minutes, washed with 50 uL of 0.5 M NaCl, and centrifuged for an additional 20 minutes. Peptide concentrations were measured with the Pierce Micro BCA assay (Thermo Scientific Pierce).

#### LC-MS/MS

We analyzed tryptic peptides with a Q Exactive HF hybrid quadrupole-Orbitrap mass spectrometer (Thermo Fisher Scientific) using a method similar to one described previously.^24^ Samples (*n*=22) collected from the Intervention cohort were run separately from those collected from the Habitual Diet participant (*n=*5). For both runs, samples were randomized and a wash run with 100% acetonitrile performed between each sample. We used an UltiMate 3000 RSLCnano Liquid Chromatograph (Thermo Fisher Scientific) to load peptides (600 ng for Intervention, 800 ng for Habitual Dietl) on a 5 mm, 300 μm ID C18 Acclaim PepMap100 pre-column (Thermo Fisher Scientific) with loading solvent A (2% acetonitrile, 0.05% TFA). Peptides were then eluted onto an EASY-Spray analytical column (75 cm× 75 µm, heated to 60 °C) packed with PepMap RSLC C18, 2 µm material (Thermo Fisher Scientific) with eluent A (0.1% formic acid in water) and eluent B (80% acetonitrile, 0.1% formic acid). We performed peptide separation using a 140 min (Intervention) or 260 min (Habitual Diet) gradient from 5-99% B at a flow rate of 300 nl/min. Peptides were ionized with electrospray ionization using the Easy-Spray source (Thermo Fisher Scientific) and mass spectra acquired in the Q Exactive HF hybrid quadrupole-Orbitrap mass spectrometer (Thermo Fisher Scientific). We performed a full MS scan from 380 to 1600 *m*/*z* at a resolution of 60,000 and maximum injection time of 200 ms. We performed data-dependent MS^2^ for the 15 most abundant ions at resolution of 15,000 and maximum injection time of 100 ms (Intervention) or 200 ms (Habitual Diet). The instrument parameters were as follows: 445.12003 lock mass, normalized collision energy equal to 24 and exclusion of ions with +1 charge state. We used a 25 s dynamic exclusion for Intervention samples and a 15 s dynamic exclusion for Habitual Diet samples. Method differences between the two sample sets are the result of LC-MS/MS method optimization.

#### Protein reference database construction

We curated a database containing: 1) the human proteome (UP000005640, Downloaded May 17, 2020), 2) microbial protein sequences derived from the human GI tract as part of the Human Microbiome Project (https://www.hmpdacc.org/hmp/HMRGD/, downloaded October 13, 2020) and 3) a custom database of protein sequences of potential dietary plants, animals, and fungi. To curate the dietary protein sequences (downloaded April 10-May 10, 2020, **Table S1**), we combined the proteomes or available protein sequences (if a complete proteome was not available) of approximately 250 different organisms that were deemed likely food items in a Durham, NC-based cohort from a global list. Every proteome (human, microbiota, individual dietary components) was independently clustered with an identity threshold of 95% using cd-hit^25^ to reduce redundancy.

A preliminary search before clustering against the most abundant human proteins revealed that some host digestive proteins (e.g. *Homo sapiens* alpha-amylase) were misidentified as dietary proteins (e.g. cattle, or *Bos taurus*, alpha-amylase). To address these misidentifications, we concatenated the individual dietary proteomes and used cd-hit-2d to remove dietary proteins with an identity threshold of at least 50% to the 15 most abundant human proteins identified in the samples (**Table S2**). This additional clustering step allowed us to be more confident that the dietary proteins we did identify were true dietary proteins and not cross-species identifications of host proteins. The final database contained 2,942,188 protein sequences.

#### Protein identification and quantification

We searched MS/MS spectra against the protein database using the Sequest HT node in Proteome Discoverer version 2.3 (Thermo Fisher Scientific) using the following parameters: trypsin (Full), maximum 2 missed cleavages, 10 ppm precursor mass tolerance, 0.1 Da fragment mass tolerance and maximum 3 equal dynamic modifications per peptide. We considered the following dynamic modifications: oxidation on M (+15.995 Da), deamidated N, Q, R (+0.984 Da), and acetyl on the protein N terminus (+42.011 Da). We also considered carbamidomethyl on C (+57.021 Da) as a static modification. We used the Percolator node in Proteome Discoverer for the peptide false discovery rate (FDR) calculation with the following parameters: maximum Delta Cn 0.05, a strict target FDR of 0.01, a relaxed target FDR of 0.05 and validation based on q-value. For protein inference, we used the Protein-FDR Validator node in Proteome Discoverer with a strict target FDR of 0.01 and a relaxed target FDR of 0.05 to restrict protein FDR to below 5%. Peptide-spectrum matches (PSMs), the taxonomic assignment of their source food, and sample metadata were organized using phyloseq v. 1.32.0.

#### Proteomic analysis

We removed a single outlier sample from the analysis that had a low number of protein identifications and peptide spectral matches compared to other samples in the dataset. Despite multiple extraction attempts, we were unable to extract a sufficient number of peptides (quantification below the limit of detection) from this sample. We chose to include the sample for LC-MS/MS analysis but omitted it from the final analysis.

Proteinaceous biomass was calculated as previously described^26^ by considering proteins with ≥2 protein unique peptides to provide high confidence that the protein originated from a specific taxon in addition to the above 5% FDR condition. Total PSMs from the remaining proteins were then summed within human, microbial taxa, and dietary taxa.

We only considered identified dietary proteins that were the master proteins of their protein group and that had at least 5 cumulative PSMs in the dataset. After summing PSMs from proteins identified to the same food taxon, we also only considered a taxon “detected” if it had ≥5 PSMs.

Because the filter we applied to the protein reference database to address misidentification of host or microbial proteins as dietary was likely incomplete, we applied an additional filter to animal-derived proteins in the dataset. We reasoned that due to a closer evolutionary relationship, animal proteins were more likely to be misidentified as human than those derived from fungi or plants. We therefore manually categorized each of these proteins with >5 PSMs as “muscle”, “egg”, “dairy” or “other” and then used regular expressions to automatically label the remainder based on the names identified in the manual pass. The regular expressions for muscle-specific proteins included matches to terms like “actin”, “titin”, “sarco-”, and “myo-”; for egg to “ovo-” and “vitello-”; and for dairy to “casein” and “butyrophilin”. In downstream analyses, we excluded the “other” category to remove potential misidentifications, though we note that this strategy also excluded uncharacterized proteins.

### Diet detection with FoodSeq

#### PCR amplification of trnL and 12SV5

*trnL* FoodSeq was performed as previously described^16^. We performed 12SV5 FoodSeq using a similar two-step PCR protocol, with the following differences in the primary amplification: 12SV5 used the AccuStart II PCR SuperMix (Quantabio, Beverly, MA) in a 10 μl volume containing 0.5 μl of 10 μM forward and reverse primers (IDT, Coralville, Iowa), 1 μl of 100 μM human blocking primer (IDT), 5 μl of 2X AccuStart SuperMix, 0.1 μl of 100X SYBR Green I (Life Technologies, Carlsbad, CA), 20 mg/μl of BSA (Thermo Fisher Scientific, Waltham, MA), 1.65 μl nuclease free water, and 1 μl of extracted DNA template. The amplification primers were 12SV5F and 12SV5R^27^ with Illumina overhang adapter sequences added at the 5′ end, and the human blocking primer was DeBarba14 HomoB^28^. Cycling conditions were an initial denaturation at 94°C for 3 minutes, followed by 35 cycles of 94°C for 20 seconds, 57°C for 15 seconds, and 72°C for 1 minute.

For both *trnL* and 12SV5, each PCR batch included a positive and negative control, and samples were only advanced to the secondary PCR if controls performed as expected (otherwise, the entire batch was repeated). Secondary PCR amplification to add Illumina adapters and dual 8-bp indices for sample multiplexing was performed in a 50 μl volume containing 5 μl of 2.5 μM forward and reverse indexing primers, 10 μl of 5X KAPA HiFi buffer, 1.5 μl of 10 mM dNTPs, 0.5 μl of 100X SYBR Green I, 0.5 μl KAPA HiFi polymerase, 22.5 μl nuclease free water, and 5 μl of primary PCR product diluted 1:100 in nuclease-free water.

#### Sequencing library preparation

Amplicons were cleaned (Ampure XP, Beckman Coulter, Brea, CA), quantified (QuantIT dsDNA assay kit, Invitrogen, Waltham, MA), and combined in equimolar ratios to create a sequencing pool. If samples could not contribute enough DNA to fully balance the pool due to low post-PCR DNA concentration, they were added up to a set volume, typically 15-20 μl. Libraries were then concentrated, gel purified, quantified by both fluorimeter and qPCR, and spiked with 30% PhiX (Illumina, San Diego, CA) to mitigate low nucleotide diversity. Paired-end sequencing was carried out on an Illumina MiniSeq system according to the manufacturer’s instructions using a 300-cycle Mid or 300-cycle High kit (Illumina, San Diego, CA, USA), depending on the number of samples in each pool. trnL and 12SV5 libraries were cleaned and pooled separately and sequenced on independent runs.

#### DNA reference database construction

A list of edible plant and animal taxa was compiled from US food availability data^29^, global surveys^30^, and reference volumes^31^. DNA sequences likely to contain *trnL* or 12SV5 were downloaded from two sources within NCBI: GenBank (all publicly available DNA sequence submissions) and the organelle genome resources of RefSeq (a curated, non-redundant subset of assembled chloroplast and mitochondrial genomes). To obtain GenBank sequences, we used the entrez_search function of rentrez v1.2.3^32^ to submit separate queries for sequences containing “trnL” in any metadata field and each plant taxon name in the Organism field (*e.g.* “*Zea mays*[ORGN] AND *trnL*" to pull data for corn, or *Z. mays*) or “12SV5” and each animal taxon name in the same manner. Sequences with an “UNVERIFIED:" flag were discarded. To obtain RefSeq sequences, the plastid and mitochondrial sequence releases current as of June 2021 were downloaded and subset to only those accessions including an edible taxon name. Results from either source were then filtered to sequences containing primer binding sites for trnL or 12SV5 primers in the correct orientation. Binding sites were identified using a custom R script with a mismatch tolerance of 20% (≤3 mismatches for trnL(UAA)g, ≤4 for trnL(UAA)h, ≤3 for 12SV5F, and ≤3 12SV5R), and sequence outside the primer binding sites removed. Identical sequences from different accessions of the same taxon were de-duplicated, but we preserved distinct sequences within taxa (indicating genetic variability) and identical sequences from different taxa (indicating genetic conservation) to yield the final references.

#### FoodSeq analysis

For each sequencing run, raw sequencing data were demultiplexed using bcl2fastq v2.20.0.422^33^. Read-through into the Illumina adapter sequence at the 3’ end was detected and right-trimmed with BBDuk v. 38.38^34^. Using cutadapt v. 3.4^35^, paired reads were filtered to those beginning with the expected primer sequence (either *trnL*(UAA)g or 12SV5F for the forward read and *trnL*(UAA)h or 12SV5R for the reverse) and then trimmed of both 5’ and 3’ sequences using a linked adapter format with a 15% error tolerance. Paired reads were quality-filtered by discarding reads with >2 expected errors and truncated at the first base with a quality score ≤2, denoised, and merged to produced amplicon sequence variants (ASVs) using DADA2 v. 1.10.0^36^. For *trnL*, taxonomic assignment was done with DADA2’s assignSpecies function, which identified ASVs by exact sequence matching to the custom trnL reference database, with multiple matches allowed. If multiple matches occurred, reads were assigned to the taxon representing the last common ancestor of all matched taxa (e.g. an ASV matching to both wheat [*Triticum aestivum*] and rye [*Secale cereale*] was relabeled as Poaceae, the family shared by both genera). For 12SV5, taxonomic assignment used DADA2’s assignTaxonomy function, as exact sequence matching had reduced performance due to the introduction of mismatches by the DNA polymerase without proofreading activity in the PCR step required for blocking primer compatibility.

Sequence data were screened for contamination on a per-PCR batch basis using decontam v1.8.0^37^ using DNA quantitation data from the library pooling step, and suspected contaminants were removed. ASV count tables, taxonomic assignments, and metadata were organized using phyloseq v1.32.0^38^.

### Dietary data collection and processing

#### Digital menus (Intervention)

Complete menu data for each participant was exported from RealChoices menu software (SciMed Solutions, Durham, NC) and linked to ingredient names from recipe source files. Ingredient common names were then manually identified to plant species using the NCBI Taxonomy Browser and Integrated Taxonomic Information System databases. For ingredients that were themselves composite foods (*e.g.,* “whole wheat bread”), we identified a primary ingredient using either provided brand information or the USDA FoodData Central database, which includes taxon mapping under the “Other information” header. For all foods, portion sizes were estimated with FoodData Central by converting the recorded menu amount (*e.g.* teaspoon, cup, ounce-weight, slice, etc.) to a gram amount using the average weights under the “Measures” header.

#### Diet records (Healthy Donor)

Dietary intake was coded from text files recorded by the participant that included descriptions of items eaten at each meal without information on quantity consumed. For ingredients that were themselves composite foods, each ingredient was coded. In cases where brand information was recorded (*e.g.* “Nutrigrain bar”, “Ben and Jerry’s Cherry Garcia”), ingredient lists were used. Otherwise (*e.g.* “pasta salad”, “sesame chicken”), ingredients were coded from a representative member of that item.

### Statistical analysis

#### Ordinations of dietary data

Diet records were binarized (assigned a value of “1” for any intake, and “0” for no intake) to synchronize data format between the Healthy Donor, who did not record quantity consumed, and the Intervention cohort, who did have estimates of quantity available. DNA- and protein-based dietary data were analyzed both as binary presence/absence and as quantity. To ordinate presence/absence data, Jaccard dissimilarity was calculated between each sample pair using the distance function of phyloseq v1.42.0 and then input into a principal coordinates analysis (PCoA) using the ordinate function of vegan v2.6.4. For quantitative data, we first computed the centered-log ratio (CLR) transform of DNA sequencing read counts or metaproteomic PSM counts to account for the compositional nature of the data using the transform function of microbiome v1.21.1. Sample-to-sample Aitchison distances (Euclidean distance between CLR-transformed counts) were then calculated on the transformed data and then input into a principal components analysis (PCA) using the function prcomp in stats v4.2.2. PERMANOVA and Mantel tests were run with the vegan functions adonis and mantel, respectively.

#### Per-food comparison between molecular measures and recorded diet

Taxon names were synchronized to account for varying detection resolution between the diet record, FoodSeq, and metaproteomic datasets. For example, menu items that were manually annotated to the subspecies or variety level (*e.g.* carrot, or *Daucus carota* subsp. *sativus*) might only be identifiable to higher levels in the molecular datasets: for instance, due to the annotation of protein sequences in public databases (*Daucus carota*) or because their FoodSeq marker region sequence is identical to other species or genera (for carrot, the same sequence is shared between carrot, parsnip, and parsley, among other species, leading the detection to be labeled as the last common ancestor of all taxa sharing that sequence, in this case the family *Apiaceae*). A complete mapping of the synchronization is provided in **Table S5**.

To assess taxon overlap between the datasets, area-proportional Euler diagrams were visualized using the euler function of package eulerR v7.0.0. Exact tests of multi-set interaction were done with function MSET of package SuperExactTest v.1.1.0. The number of taxa detected within each sample was compared with repeated-measures ANOVA using the lm function of stats v4.2.2 and Anova of car v3.1.2.

We compared DNA- or protein-based presence or absence to the presence or absence of the same food taxon in the menu record from 1 to 2 days prior to account for the mean (28 h) and typical variation of measured gastrointestinal transit times in humans (28, 29). Responses were coded as true positives (TP, food present by both molecular detection and menu), true negatives (TN, absent by both molecular detection and menu), false positives (FP, present by molecular detection, not by menu), and false negatives (FN, absent by molecular detection, but present in menu). Recall was calculated as TP/(TP+FN). Precision was calculated as TP/(TP+FP). Two-tailed Spearman correlations between precision and recall were performed using the cor.test function from R stats v4.1.3.

### Data availability

The metaproteomics data and protein sequence database have been deposited to the PRIDE repository^39^ with the dataset identifiers PXD029334 [Reviewer Access at https://www.ebi.ac.uk/pride/login User: Password: TwNtwIgz] for the participant on the habitual diet and PXD030004 [User: Password: eRGshuIR] for the diet intervention participants. FoodSeq data will be uploaded to ENA prior to publication.

## Results

### DNA- and protein-based detection of consumed foods

We conducted paired assessment of food-derived proteins and DNA in stool collected by five individuals: four volunteers consuming interventional diets at a medically supervised weight loss center, and one donor consuming their typical diet (“Intervention” and “Habitual Diet”, respectively, *n*=27 total samples). The samples shared two features that enabled the joint evaluation of metaproteomics and metabarcoding (FoodSeq): (1) they were accompanied by detailed, longitudinal dietary records for external validation (**Fig. 1a**), and (2) the majority were collected in runs of several consecutive days to investigate the kinetics of molecular dietary signals. Measurement of peptide spectra by metaproteomics, amplification of the *trnL*-P6 chloroplast marker region (“plant FoodSeq”), and amplification of the V5 region of the mitochondrial 12S rRNA gene (“animal FoodSeq”) was successful for 26, 23, and 27 of the samples, respectively. In successful samples, we detected a median of 1,985 metaproteomic peptide-spectrum matches (range 794-17,472), 68,256 *trnL* reads (range 9,744-95,249), and 7,647 12SV5 reads (range 404-22,705; **Fig. S1**. To identify foods by their peptide spectra or DNA sequences, we generated reference databases of protein or marker gene sequences from a manually curated list of 246 foods known to be consumed in human diets. The protein reference database included proteomes from 180 plants, 56 animals, and 8 fungi (**Table S1**) and was refined to exclude animal proteins similar to the human proteins that are most abundant in human fecal material (**Table S2**). The DNA reference database contained 909 sequences representing 591 plants and 31 animals. Collectively, peptides from 8,273 unique food-derived proteins and 93 DNA amplicon sequence variants (ASVs; 82 [88%] from *trnL*, and 11 [12%] from 12SV5) were detected in the sample set.

**Figure 1.**
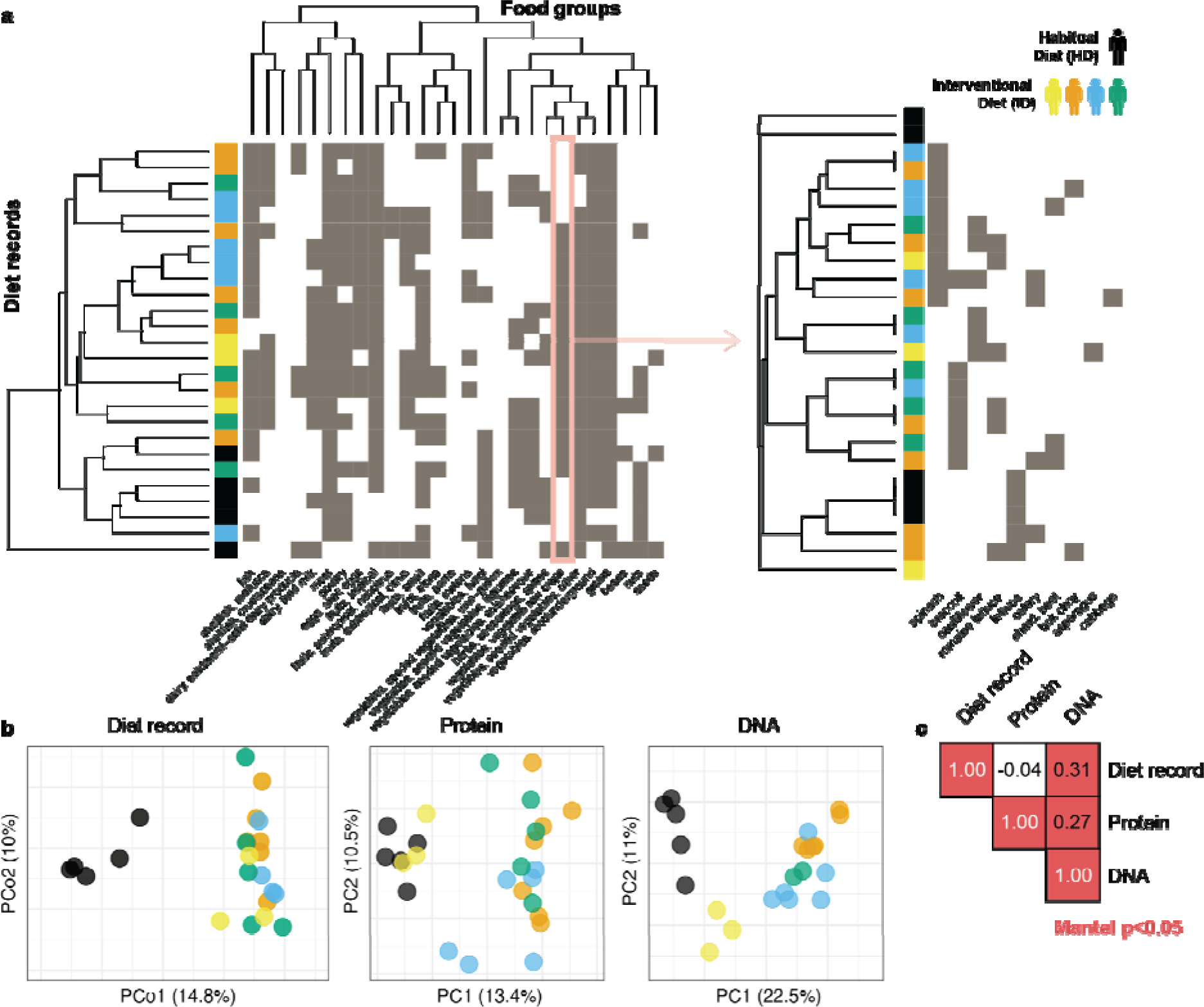
Dietary landscapes of study participants by written records and stool-based measurements. (**a**) Participant diets included food items derived from 27 food groups, shown as a heatmap of presence (gray) or absence (white) of each food group (x-axis) in the recorded diet on the day prior to stool sampling (y-axis). The x-axis dendrogram reflects food group relationships as structured by the Interagency Food Safety Analytics Collaboration (IFSAC)^40^ and the y-axis dendrogram the clustering of recorded diets by relatedness of the food groups they contain. Although food groups are displayed for clarity, dietary records provided data resolved to the level of individual food species (*e.g.* the column “vegetables, vegetable row crops” summarizes data on 13 unique food items, shown in inset). (**b**) Principal coordinate analysis (PCoA) and principal component analysis (PCA) ordinations of samples in dietary space derived from either recorded diet, metaproteomic detection of food proteins in stool, or FoodSeq detection of DNA in stool. Points represent individual stool samples, or for menu data, the day of eating prior to sample collection. Samples are colored by participant and by diet type, either habitual diet (HD) or interventional diet (ID1 to ID4). Metaproteomic data are filtered with the final criteria described in the text (5% FDR, >=1 UP and >5 PSMs for the food taxon). (**c**) Results of Mantel tests comparing distance matrices of points in **(b)** between datasets, interpretable as correlation coefficients (1, perfect positive correlation; 0, uncorrelated; -1, perfect negative correlation).

In the metaproteomic dataset, proteinaceous biomass was dominated by microbiota-derived proteins (74% of all peptide-spectrum matches), with smaller contributions from host (14%) and dietar (11%) proteins (**Fig. S2**). To tune the sensitivity and stringency of the metaproteomic analysis for dietar proteins, we tested two filtering methods: (1) selecting dietary proteins identified with a 5% false-discovery rate (FDR) cutoff and at least one *protein* unique peptide and (2) selecting dietary proteins identified with an FDR of 5% and at least one *protein group* unique peptide. The first analysis restricted proteins to those definitively identified by one or more peptides matching to a single protein sequence, whereas the second included proteins that did not have a unique peptide match, but for which there was a unique peptide match to the protein group composed of very similar homologous sequences. Because we observed only slight variations in the overall results between the protein-unique peptide (PUP) strategy and the unique peptide (UP) analysis and the UP analysis included 39% more PSMs, 2,616 additional proteins, and 18 additional food taxa (**Fig. S3**), we selected the UP analysis for all downstream steps.

Despite the steps taken to remove dietary proteins similar to human proteins from the metaproteomic reference database, we noted detection of a high number of food species (*n*=18) in every sample, including staple foods like wheat, corn, and soy, but also less common items like goat and salmon. We also observed persistent detection of digestive tract and intestinal epithelial proteins (*e.g.* progastricin, enteropeptidase, glycoprotein 2). We therefore categorized every protein name from an animal species with ≥5 cumulative PSMs in the full dataset as “muscle”, “egg”, “dairy”, or “other” (n=280 manual categorizations) and used regular expressions to automatically label the remainder (animal-derived proteins with <5 cumulative PSMs, n=1,170) with the same dietary categories (see **Table S3**). In downstream analyses of animal taxa, we excluded the “other” category, which included likely host and microbial cross-identifications as well as uncharacterized proteins. When considering food taxa identified by metaproteomics generally, we filtered to only those taxa with five or more PSMs (46 taxa removed); we performed no additional filtering when analyzing data at the level of individual proteins. Across samples, 105 foods were detected by metaproteomics with the most abundant ones being rice (*Oryza sativa*), oats (*Avena sativa*), and chicken (*Gallus gallus*); out of 90 foods detected by FoodSeq, the most prevalent items were carrot family (a DNA sequence variant shared by carrots, celery, parsley, and parsnips), peppers, avocado, cattle, chicken, and turkey (**Table S4**).

### Overall dietary assessment by record, DNA, and protein

Because there is no gold-standard method for dietary assessment, we first evaluated the consistency between FoodSeq, metaproteomics, and conventional diet records as tools to capture inter- and intra-individual dietary differences. Participant identity was significantly associated with the overall composition of the diet provided by all three techniques (**Fig. 1b**). Roughly 22% of the variation in dietary composition captured in menu records could be attributed to differences between the participants (*PERMANOVA* on Jaccard dissimilarity, *R^2^*=0.22, p=0.001). This individual-specific dietary signature was also present in both food DNA and protein composition in stool, and was stronger for DNA-based than protein-based data (*PERMANOVA* on Aitchison distance *R^2^*=0.45 for FoodSeq, and 0.28 for metaproteomics, both with p=0.001; **Fig. 1b**).

We tested if sample-to-sample differences were similar between the three measures of diet composition using the Mantel test, which evaluates correlation between distance matrices. When we used distance metrics that incorporated abundance data (Aitchison distance on the number of metaproteomic peptide-spectrum matches or DNA sequencing read counts), inter-sample distances were significantly and positively correlated for DNA and protein (Mantel *r=*0.265, p=0.001), DNA and menu (*r=*0.307, p=0.002), but not for menu and protein (*r=*-0.0453, p=0.659; all visualized in **Fig. 1c**). Interestingly, only the relationship between menu- and DNA-based diet composition was preserved when we used a presence/absence-based distance, indicating potentially significant quantitative information present in the molecular datasets (Jaccard dissimilarity, **Fig. S4**.

### Direct comparison of molecular detection to recorded diet

We next evaluated FoodSeq and metaproteomic data in relationship to recorded dietary data. In doing so, we note that neither cohort’s dietary data represented a “ground truth” comparator. The manually recorded diet of the healthy donor lacked information on amount of food and exact ingredients of prepared dishes that were present in the detailed digital menus from the Intervention center; the Intervention menus reported only food served (rather than consumed) and did not track items that could be freely chosen (*e.g* fresh fruit and salad bar offerings that varied daily and potential off-menu eating that occurred when clients were not at the center). However, exact menu records or diet diaries are established methods of dietary assessment, so we evaluated molecular measures against them.

All food items were coded to a plant, animal, or fungal source taxon. We manually synchronized taxon names across the three datasets to reconcile naming differences that arose by data type (see **Methods**, **Table S5**. There was significant overlap between taxa identified by the three datasets, with the size of the intersection of taxa observed by all three measures unlikely to be detected by chance (multi-set intersection test p=0.004 for plant and p=0.014 for animal taxa, **Fig. 2a**). 66% of plant taxa and 38% of animal taxa were detected by at least two measures. Comparing the number of food taxa identified within single samples, there was no difference between the number of plants recorded in the menu and detected by *trnL* FoodSeq (repeated-measures ANOVA, Benjamini-Hochberg adjusted p=0.31); however, 12SV5 identified significantly fewer animal taxa (p=0.002) and metaproteomics identified significantly more plant and animal taxa within each individual sample (p<10^-12^ and p=0.002 for plant and animal taxa, respectively, in comparison to menu; **Fig. 2b**).

**Figure 2.**
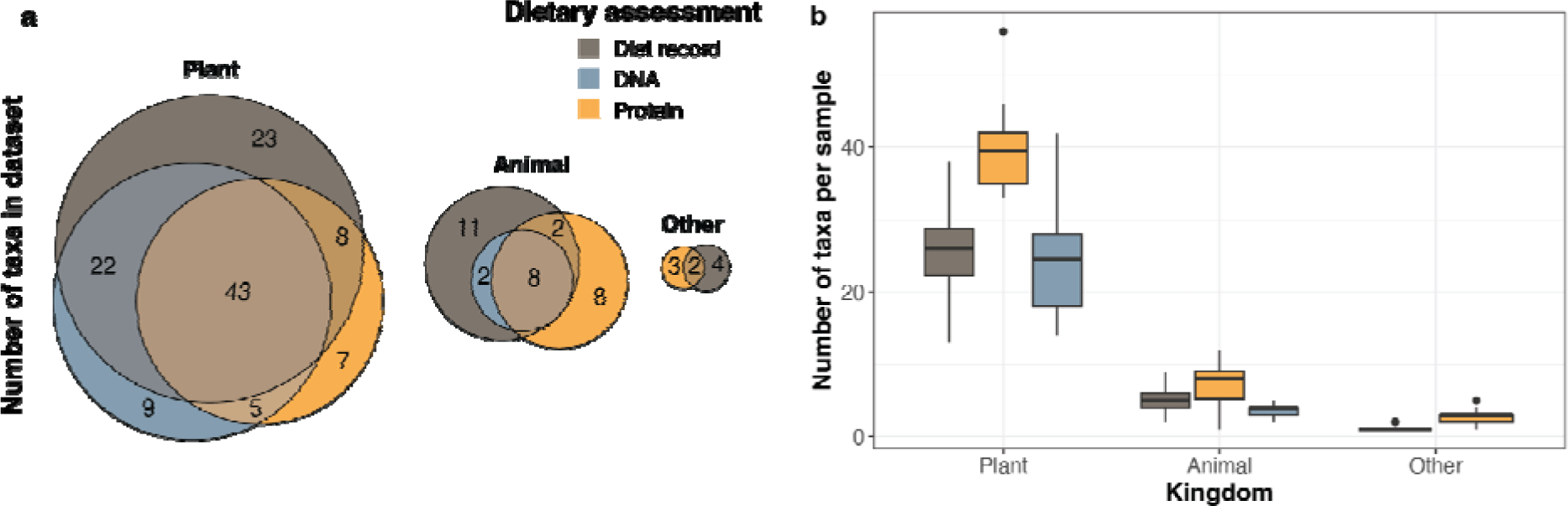
Diet records and stool measurements include similar food taxa. **(a)** Overlap between food taxa detected by diet record, stool DNA, or stool protein, separated by kingdom. Note that the FoodSeq assay does not include a marker for fungi, seaweed, or bacteria-derived foods (*i.e.* xanthan gum), which are shown in the “Other” category. For this analysis, menu data is included from all available records, which in some cases may be up to 4 days from the closest sample. Molecular data is only included from samples with successful molecular detection by both protein- and DNA-based methods, which is why the total number of taxa differs slightly from that in all samples, reported in **Table S4**. **(b)** On a per-sample basis, estimates of richness (the number of unique food items recorded or detected) were significantly higher by metaproteomics than for the two other measures.

In accordance with estimates of typical gastrointestinal transit time at 24-48 hours (28, 29), we calculated FoodSeq and metaproteomic performance in comparison to the prior two days of recorded diet at the level of food taxon. Cumulatively, performance was higher for FoodSeq than metaproteomics across taxonomic ranks (**Fig. 3a**; two-way ANOVA p<10^-5^ for performance by dataset, p=0.4 for performance by taxonomic level) and, within each measure, significantly higher for animal compared to plant taxa (unpaired Mann-Whitney test, p<10^-5^ and p=0.005 for DNA and protein, respectively; **Fig. 3b**). Per-taxon performance varied widely, with some taxa in near-perfect agreement with menu data and others dominated by false positives, false negatives, or a mixture (**Fig. 3c**).

**Figure 3.**
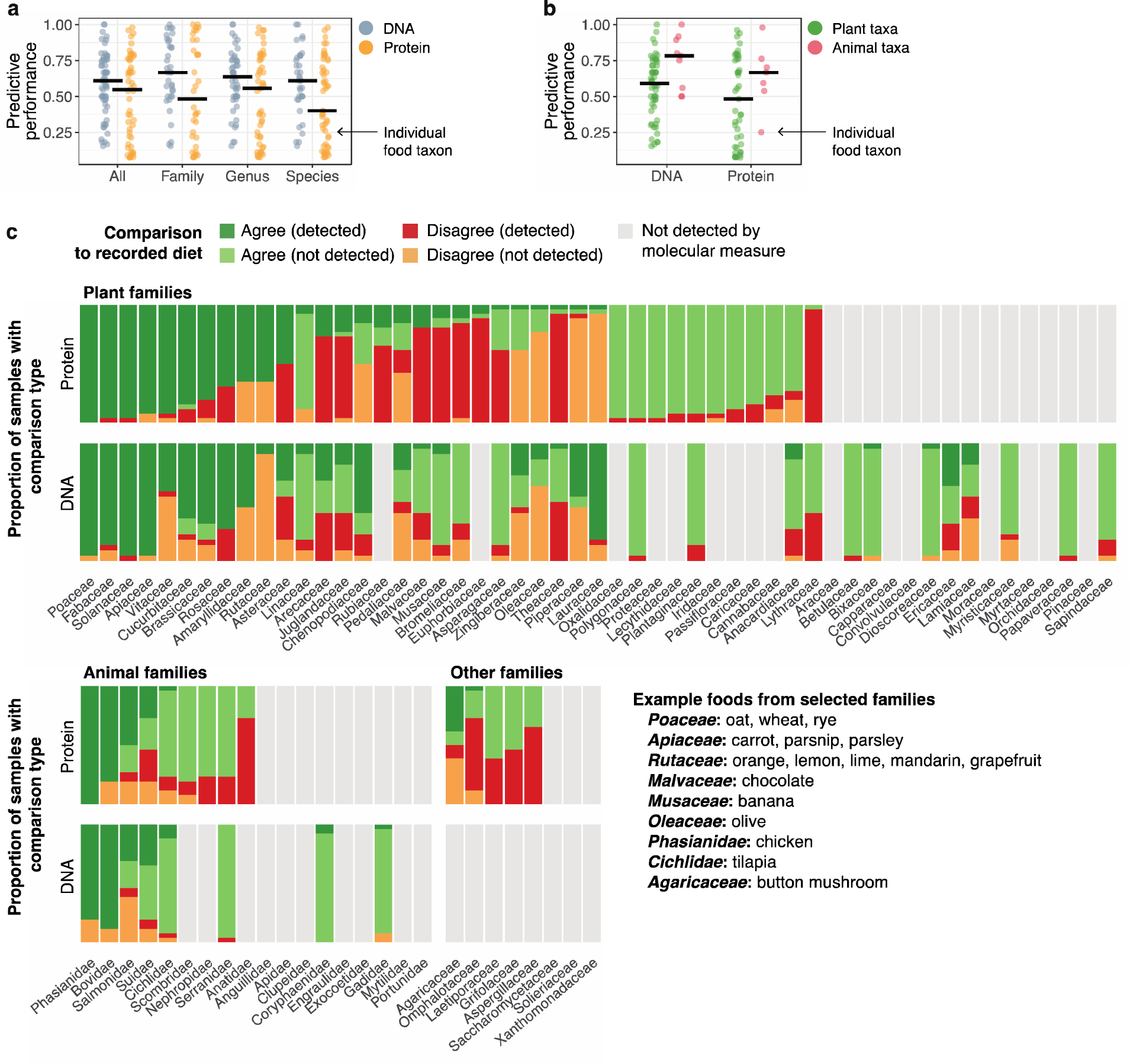
Performance of DNA- and protein-based dietary assessments in comparison to recorded diet. **(a)** Summary of molecular detection consistency with menu data across taxonomic levels of analysis. “All” levels indicates no aggregation of taxa, preserving each individual taxon at the level to which it can most accurately be specified by the three methods (a mix of family, genus, and species designations). The “predictive performance” measure used here is the F-measure, which is the harmonic mean of precision and recall and ranges from 0 (completely inaccurate detection) to 1 (perfect precision and recall). Black bars indicate the median. **(b)** Comparison of performance by DNA- and protein-based assessment in comparison to recorded diet between food taxa of animal and plant origin. **(c)** Protein and DNA detections in comparison to the recorded diet from the two days prior to sample collection. For ease of visualization, data are presented at the family level; see **Fig. S5** for a visualization of the full dataset. Detections were coded as true positives (TP, food present by both molecular measure and menu), true negatives (TN, absent by both molecular measure and menu), false positives (FP, present by molecular measure, not by menu), and false negatives (FN, absent by molecular measure, but present in menu). Taxa are aggregated to the family level and ranked for each method in descending magnitude of its F measure statistic for their detection, which is the harmonic mean of sensitivity and positive predictive value).

Because of the limitations of the menu data noted above, we also directly compared FoodSeq and metaproteomic performance to determine if divergences from menu data were shared or distinct. For each measure, we calculated recall (or true positive rate, the proportion of recorded foods that were detected by molecular signal) and precision (or positive predictive value, the proportion of positive molecular signals that were confirmed by a menu entry). On a per-taxon basis, FoodSeq and metaproteomic recall were moderately correlated with one another (Spearman 17=0.02-0.52, p∼10^-1^-10^-4^ across varying taxonomic rank), but for precision this correlation was markedly stronger (Spearman 17=0.80-0.88, p∼10^-11^-10^-14^). This finding was consistent across taxonomic levels (**Fig. 4**). Recall and precision differ by only one term in their denominator: the number of false negatives (recall) or the number of false positives (precision). The increased correlation observed for precision indicated that the false positive structure between the two molecular measures was stronger than the false negative structure. In other words, a false positive by either measure was more likely to be shared by the alternate approach, and could be reflective of a real divergence from the recorded menu (a forgotten food, food eaten off menu, or an error in assuming a constant two day lag in our analysis). However, failures to detect recorded items were less correlated with one another, and therefore likely to be due to method-specific limitations or biases. From this comparison, we also noted extremes of detection: some foods were reliably detected by both measures (chicken, peppers, and carrots), better detected by FoodSeq (tilapia, peas), better detected by metaproteomics (citrus), and poorly suited for detection by either method (olives or their oil, tea).

**Figure 4.**
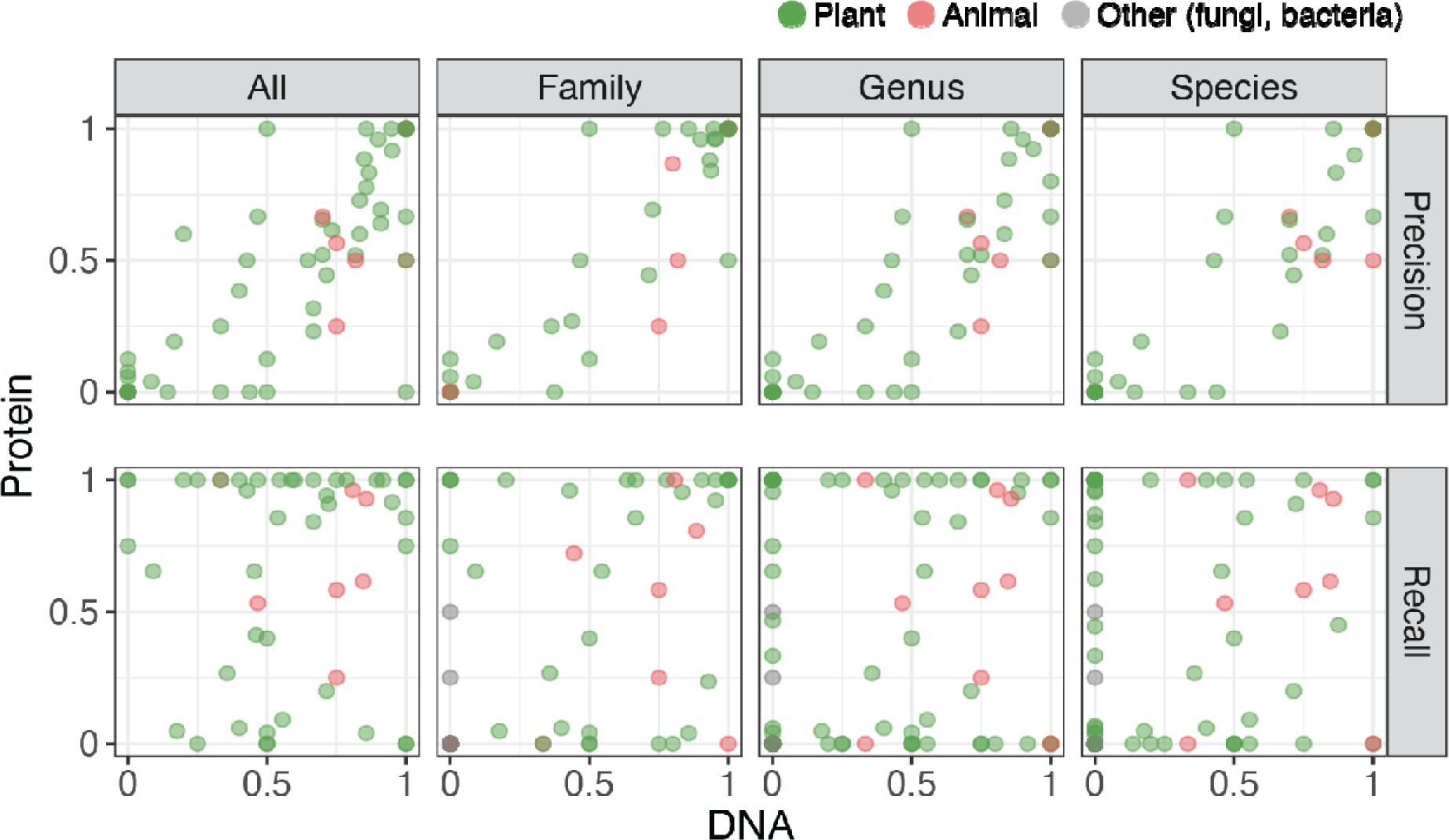
Per-food detections by metaproteomics and FoodSeq are more correlated for precision than recall. Each point represents an individual food taxon, summarized to the taxonomic level as labeled at the top of each facet. “All” includes every taxon in the dataset without any aggregation; these can differ in their taxonomic level due to a variable degree of resolution in the FoodSeq data.

#### Metaproteomics enables distinction of food type from the same source species

Finally, we selected a subset of foods for analysis to highlight the tissue-specificity of metaproteomics and support its use as a complementary measure to FoodSeq. Unlike DNA signatures, which are common across all tissues, metaproteomics provides data on individual protein constituent that reflect tissue of origin and thus food type. We began by specifically investigating *Gallus gallus* (chicken) and *Bos taurus* (cattle), which are food species with prominent tissue-specific consumption of meat versus egg or dairy. In the metaproteomic dataset, we labeled 28, 10, and 4 proteins as meat- specific, egg-specific, and dairy-specific, respectively (**Table S3**) and used them to categorize intake from stool. The key proteins identified for meat were myosin, actin, and titin; for egg, avidin, ovotransferrin, ovalbumin (abundant in egg white), and vitellogenins (abundant in yolks); and for dairy, caseins, butyrophilin (the major protein associated with fat droplets), and beta-lactoglobulin (the major whey protein in cow’s milk). For all tissue types but dairy, detections by tissue-specific proteins had no significant difference from presence or absence of the same food tissue in diet records from the day prior to sampling (McNemar test p=1.0 for egg, p=0.7 for chicken meat, p=0.7 for cattle meat, and p=0.0008 for dairy; **Figure 5**). Metaproteomics detected dairy intake less frequently than menu records (dairy detected in 50% of samples compared to 100% of the diet records one or two days prior to sampling), potentiall reflecting the impact of processing, digestion, or lower protein content per gram of tissue.

**Figure 5.**
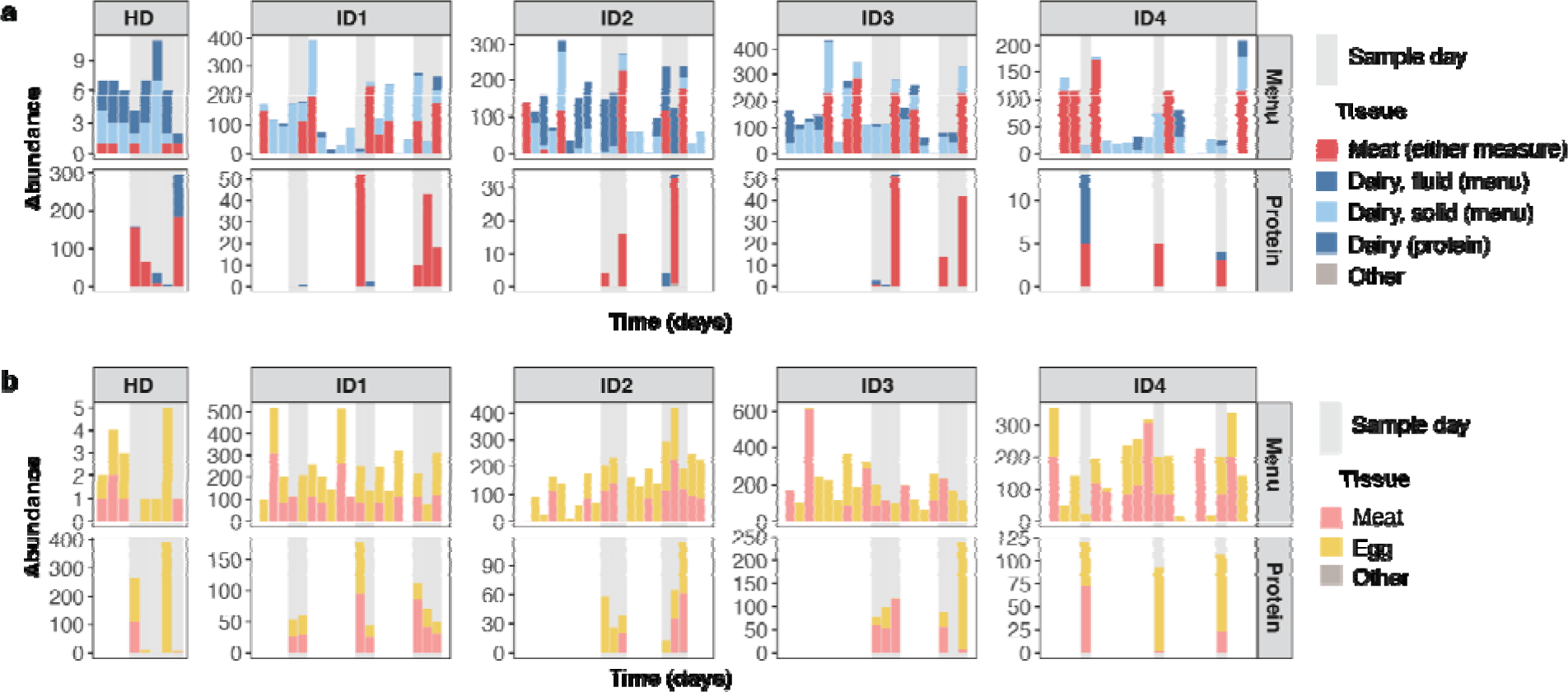
Metaproteomics enables food type-specific detection. Menu records (upper panel) and metaproteomic detections (lower panel) of food type-specific intake of cattle **(a)** and chicken **(b)** intake. Dates within each subject are relative, days on which a sample was collected are indicated in gray, and the menu and protein panels are not shifted relative to one another in time. A protein detection should therefore be compared to the several preceding days of recorded intake in the menu panel. Dietary abundance in the habitual diet participant (HD) was recorded as instances of intake (for example, 1 for consumed once in that day, 2 for consumed twice, etc.), and in the interventional participants as grams estimated from ordered servings of individual recipes.

We next tested if we could identify a set of candidate protein biomarkers for specific foods in the metaproteomic dataset that would lend themselves to future development of targeted proteomic assays for high throughput detection at reduced cost. Considering foods detected in ≥5 samples (n=85), we identified individual proteins that were present in more than half the cases an individual food taxon was detected by metaproteomics. Of 3,392 candidate proteins for the initial 85 foods, we identified 67 candidates with this strategy that could serve as potential standalone indicators of intake, covering food species including corn, oats, chocolate, brussels sprouts, almond, grape, and chicken (**Table S6**). In addition, we also noted interesting cases that did not meet these criteria (potentially due to mixing of true signal with spurious positive detections from food proteins that share similarity to human proteins). Two specific additional proteins we want to highlight as examples are ananain and bromelain, which are both proteases that are highly specific to pineapple. Although daily fruit offerings in Intervention participant were not always recorded to the species level, three intake events for bromelain came shortly after recorded meals with pineapple as an ingredient in this cohort and ananain was detected in the Habitual Diet participant only in the sample collected the day after written pineapple consumption.

## Discussion

We have shown that residual dietary DNA and protein from consumed foods are present in human stool, readily detected with existing techniques, and can be identified with reference database curated from publicly available genomes of food species. Our findings demonstrate that both FoodSeq and metaproteomics detect a broad range of foods and have a significantly similar global structure. Thi detection is possible despite forces that may degrade the molecular components of food prior to detection, including thermal and chemical degradation by cooking and enzymatic degradation in the gastrointestinal tract by host proteases and nucleases. In our data, dietary sources still accounted for a substantial proportion of total proteinaceous biomass of stool (∼11% on average). Comparable data on the prevalence of dietary DNA in the overall pool of extracted stool DNA is sparse but supportive of a lesser contribution: in a wild primate model, 0.004–0.008% of metagenomic reads were dietary in origin,^41^ which agrees with anecdotal communications we have heard from researchers working in humans (unidentified metagenomic sequences assumed to be dietary and <1% of total reads). An increased prevalence of diet-derived stool protein in comparison to DNA also agrees with prior work in invertebrate animals showing that protein epitopes, especially those internal to the protein, survive in the gut for longer than DNA fragments.^17^

Despite likely lower input, we found that on average, DNA-based assessment had higher detection performance than protein when we considered individual foods in direct comparison to the diet record (**Fig. 3a**). FoodSeq protocols include an amplification step that metaproteomics does not, potentially allowing DNA to more reliably report food items present in lower initial abundance like herbs and spices. This would lead to false negative detections by protein (protein failing to detect a recorded item) in the context of true positive detections by DNA. However, we noted that false positive, not false negative, detections were the most common discrepancy between protein and menu data. For the animal food taxa we suspect that in some cases, this may be due to continued misidentification of host-derived proteins as dietary ones by our necessarily large reference database. For the plant taxa these “false positives” are likely due to a mix of reasons including (1) fruit options consumed by participants that were not recorded on the menu such as pineapple (Bromeliaceae), (2) foods consumed by participants off menu such as products containing chocolate (Malvaceae), and (3) the incompleteness of some reference food genomes in public databases, which can lead to mis-identification of close relatives.

23 food taxa were detected in >90% of samples by metaproteomics: these included several species that were supported by diet records and could plausibly be consumed daily by the US-based participants in this study (corn, chicken, soy) but others that were not (dates, Napa cabbage, cassava). We were able to reduce putative and false positive signals from prevalent animal detections (goat, duck) by considering only animal-derived proteins that we could definitively identify as originating from skeletal muscle (meat), egg, or milk (dairy). However, we did not perform a similar filtering strategy for plant foods due to the higher variation in tissue types consumed (*e.g.* roots, tubers, leaves, fruits, seeds, nuts, grains, etc.). One of the potential reasons for the incorrect identification of plant taxa is likely the fact that many of the protein sequence databases for plant derived foods are still incomplete. If sufficient similarity between protein sequences is present, proteins from a species with an incomplete proteome may instead be identified as one or more related taxa. For example, very few protein sequences are available for broccoli (*Brassica oleracea* var. italica) at the moment. Therefore, if a participant consumed high quantities of broccoli and thus many mass spectra of broccoli peptides are acquired from stool samples, these spectra will likely match to proteins from other members of the cabbage family (Brassicaceae) of which broccoli is a member. Future sequencing efforts of food species genomes should improve database quality and resolve associated taxonomic resolution issues. Additionally, if full genomes cannot be acquired, the sequencing of specific genes from food species could be used to obtain the sequences of dominant food proteins to be able to discriminate food species. The false positives caused by database incompleteness may also have contributed to the overall weaker correspondence we observed between proteomic data globally and dietary records by Mantel test.

Nevertheless, proteomic data can provide more dietary information than DNA alone. DNA cannot distinguish foods with an identical sequence at the marker region used: this is the case not only for foods derived from different parts of the same organism but often for foods derived from closely related species. For instance, a single *trnL* sequence variant is shared by dill, carrot, cumin, parsley, fennel, and parsnip, but these foods are readily distinguished by protein constituents in our data. Tissue type conveys important nutritional information (*e.g.* the fat content of chicken breast versus egg) that is unmeasurable by FoodSeq but readily assessed by metaproteomics. In our data, we identified multiple cases where metaproteomics had higher resolution than FoodSeq. Protein signals differentiated durum wheat (most commonly used in pasta) from bread wheat, and identified distinct tissues from the same food species (*e.g.* chicken and egg, beef and dairy). For future biomarker development, food-derived proteins provide a large number of candidate targets, some of which are very abundant in specific tissues. Additionally, the detection limit of protein-based biomarkers could be increased by developing targeted mass spectrometry approaches that are much more sensitive than the discovery-focused untargeted approach that we used.^42^

Features of the consumed diet and its record limited our analyses, and we can therefore make specific recommendations for the design of future dietary studies and follow-on validation. To assess the performance of FoodSeq and metaproteomics in real-world diets and across many foods, we included participants with diets that included a range of items and varied day-to-day. However, in some cases, this limited our analysis: for example, most participants consumed both beef and dairy or both chicken and egg in the days prior to sampling, which precluded a clear connection between detected proteins and a specific episode of prior intake. Designing diet interventions that include non-repeating food items or introduction of a specific item into the baseline diet may help to adjust for this effect by more accurately linking molecular and menu data. Even though the dietary records of both Intervention and Habitual Diet participants were detailed, they had notable limitations: some foods were not tracked by the digital menu system, and others could be eaten off-menu or left on the plate (Intervention); quantity was not recorded, and complex meals were not always separated into their component ingredients (Habitual Diet). Future studies could include paired sampling of food inputs and stool outputs in order to reduce errors from ingredients inferred during the process of coding written records to food items.

The data shared here can be re-analyzed for food-specific candidate biomarkers that would lend themselves to development in targeted proteomic assays that would allow for high throughput and reduced cost. Clear frameworks for establishing biomarker validity from nutritional epidemiology^43^ can then guide *in vitro* and *in vivo* testing. With additional development, metaproteomics may be helpful not only for detecting what is present in stool, but to determine what is *absent* from foods known to be consumed, and thus being degraded farther upstream and absorbed by the host or fermented by microbiota. A central theme in dietary assessment has been developing tools with different sources of error that can be used to validate one another. By expanding the range of tools available for dietary assessment, our work provides independent measures of food and tissue constituents in the diet that can be used in future studies.

## Acknowledgments

We thank our study volunteers for their participation, Verónica Palacios for human study support, and Tonya Snipes, Lisa Alston-Latta, and Margaret Huggins for keeping our lab spaces and glassware clean. We thank Angie Mordant and Clara Tang for assistance in metaproteomics data generation. This work was supported by the National Institute Of General Medical Sciences of the National Institutes of Health under Award Numbers R35GM138362 (MK) and R01DK116187 (LAD), the Foundation for Food and Agriculture Research Grant ID: 593607 (MK), the Duke Microbiome Center (LAD), the Triangle Center for Evolutionary Medicine (BLP), the Integrative Bioinformatics for Investigating and Engineering Microbiomes Graduate Student Fellowship (BLP), and the Ruth L. Kirschstein National Research Service Award to the Duke Medical Scientist Training Program. This work used a high-performance computing facility partially supported by grants 2016-IDG-1013 (“HARDAC+: Reproducible HPC for Next-generation Genomics”) and 2020-IIG-2109 (“HARDAC-M: Enabling memory-intensive computation for genomics”) from the North Carolina Biotechnology Center. We made all LC-MS/MS measurements in the Molecular Education, Technology, and Research Innovation Center (METRIC) at North Carolina State University.

## Extended data

**Figure S1.**
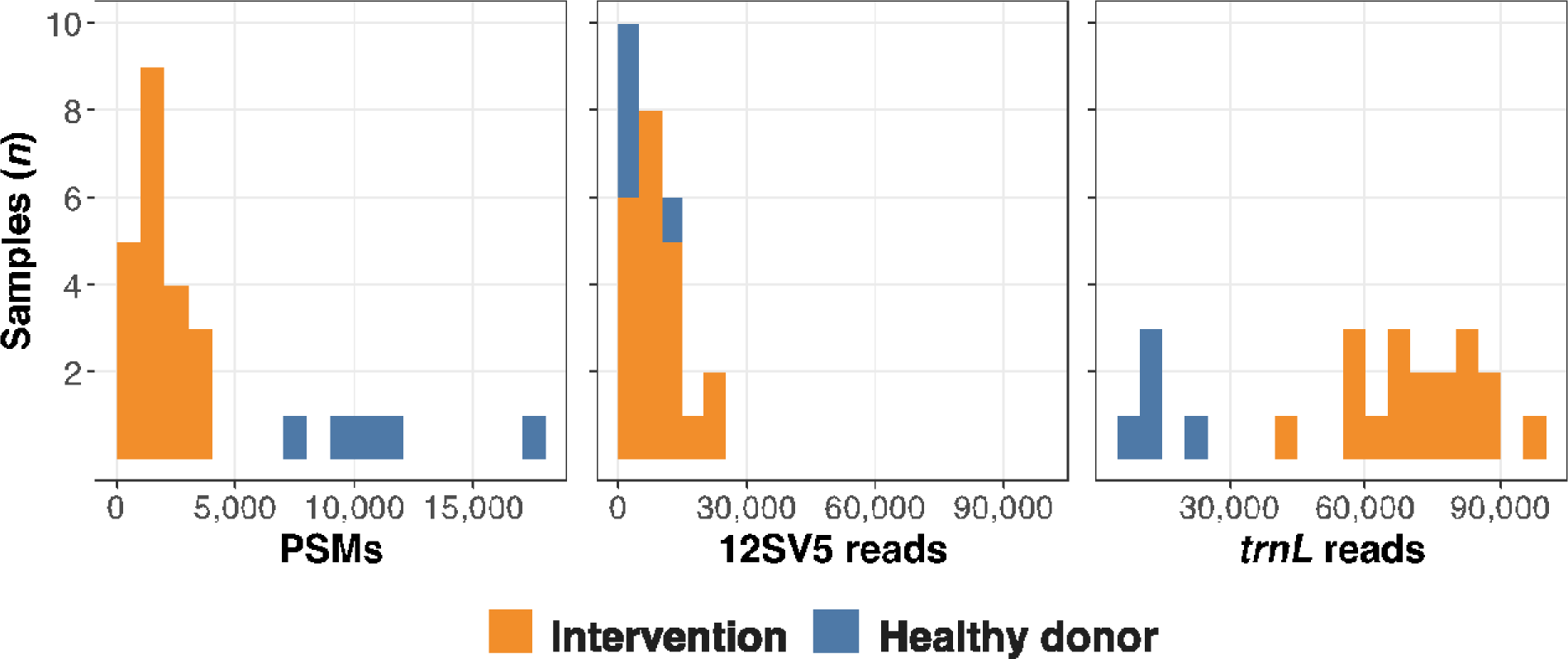
Detection of food-derived protein and DNA by sample and cohort. Sampling depths of peptide-spectrum matches (PSMs) and FoodSeq reads in stool samples by cohort. For metaproteomics, healthy donor samples were run with a longer LC gradient (140 compared to 260 minutes), leading to higher PSMs/sample. FoodSeq samples were processed in batches by study and marker, leading to variations in read depth due to overall run metrics (notable for *trnL*).

**Figure S2.**
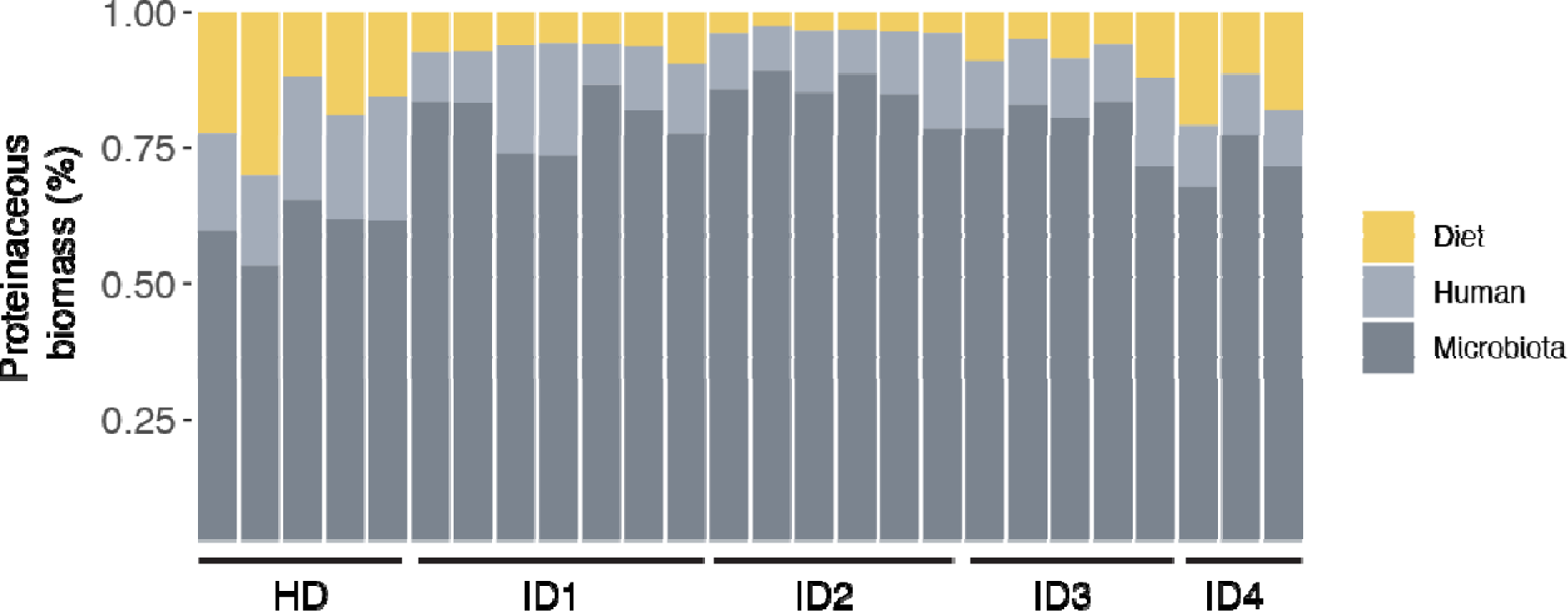
Proteinaceous biomass by sample. Bars indicate individual samples and are grouped by subject (HD, Healthy Donor; WL1-4, Intervention subjects 1 to 4).

**Figure S3.**
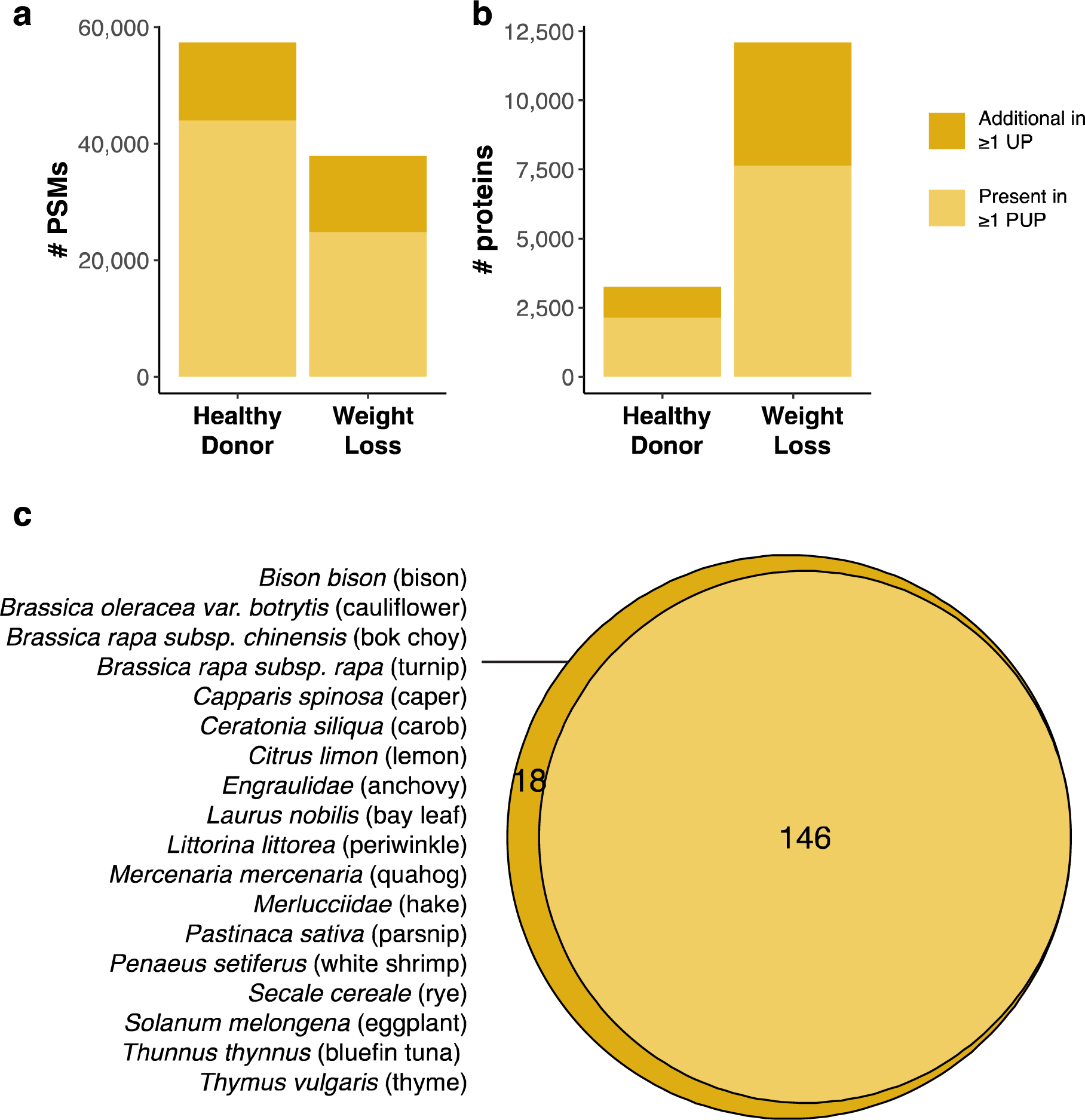
Comparison of filtering strategies for metaproteomic data. Number of **(a)** peptide-spectrum matches (PSMs), **(b)** proteins, and **(c)** food taxa detected by filtering proteins for either ≥1 protein unique peptide (≥1 PUP) or ≥1 unique peptide (≥1 UP).

**Figure S4.**
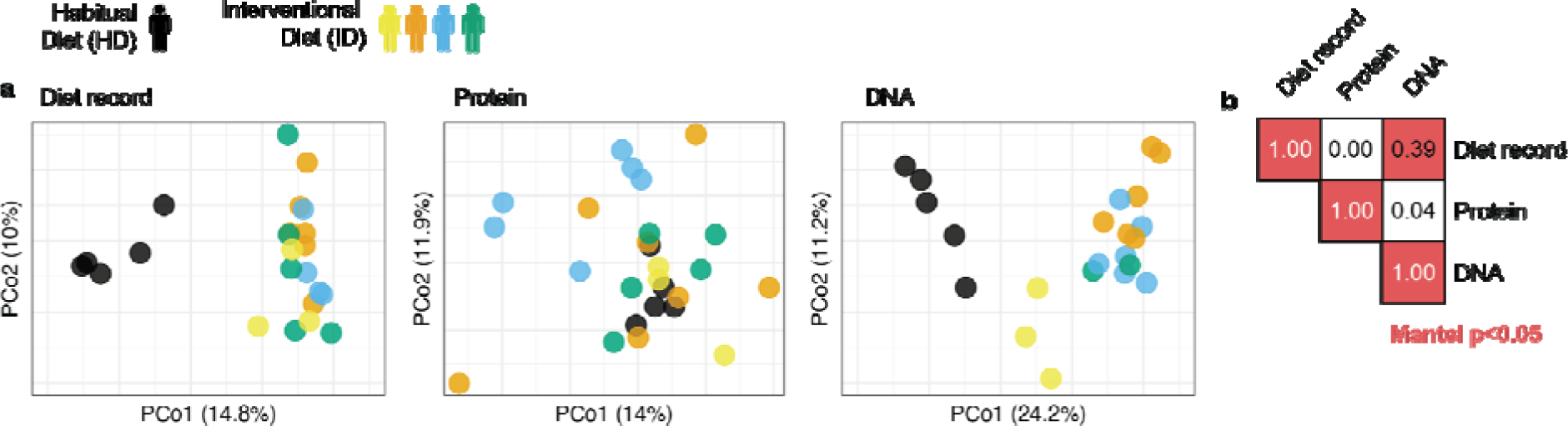
Only DNA-to-diet record associations in overall diet composition are preserved in a presence-absence framework. Analysis as in **Fig. 1b,c** with principal coordinate ordinations for all measures and Jaccard dissimilarity as input to Mantel tests.

**Figure S5.**
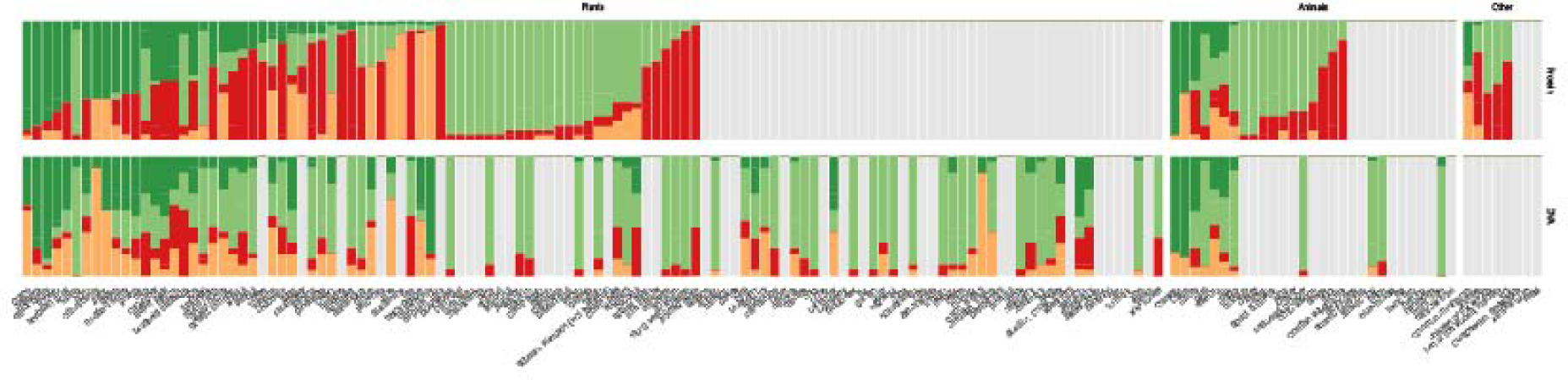
Performance of DNA- and protein-based dietary assessments in comparison to recorded diet across all taxa. Because FoodSeq, metaproteomics, and written records can specify foods with different levels of precision, comparisons between the measures do not always occur at a set taxonomic rank (*e.g.* species). Shown here is a version of Fig. 3, which showed performance at the family level for succinctness in the main text, that now includes all detected or recorded food items by any measure to the most precise taxonomic level that they can be identified in synchrony by the three methods. To do so, taxon names were manually matched across DNA, protein, and menu data to create a mapping between the three measures (**Table S5**). This also allows us to plot the data using the common names for foods, which are not always easily mapped onto family, genus, or species labels. This representation of the data is what is summarized in the “All” category shown in **Figs. 3a,b** and **4**. More details are described in **Methods**.

